# Structural framework for the assembly of the human tRNA ligase complex

**DOI:** 10.1101/2025.08.01.668197

**Authors:** Moritz M Pfleiderer, Moritz Leitner, Ajse S. Nievergelt, Alena Kroupova, Javier Martinez, Martin Jinek

## Abstract

In human cells, a subset of tRNA-encoding genes contain introns. These are removed by a non-canonical splicing pathway in which the tRNA splicing endonuclease complex catalyzes intron excision and the resulting exons are subsequently ligated by the tRNA-ligase complex (tRNA-LC). Although recent studies have provided insights into the process of intron removal, the molecular mechanisms underpinning tRNA ligation by tRNA-LC remain elusive. The tRNA-LC is a hetero-pentameric protein assembly consisting of Ashwin, CGI-99, FAM98B, the DEAD-box helicase DDX1 and the catalytic subunit RTCB/HSPC117. Using cryo-EM, we have determined an atomic-resolution reconstruction of human tRNA-LC. We find that CGI-99, DDX1 and FAM98B form an alpha-helical bundle that contacts RTCB via an interface located on the opposite side from the location of the ligase active site and tethers DDX1 to the tRNA-LC via its C-terminal helix. FAM98B and CGI-99 extensively interact in an intricately co-folded heterodimer that clamps Ashwin in a pincer-like structure. Interaction analysis using structure-based mutants of tRNA-LC subunits supports the overall architecture of the complex. Finally, we show that the paralogous proteins FAM98A and FAM98C underpin the assembly of compositionally distinct RTCB-containing complexes that lack Ashwin and may have distinct cellular functions. Together, our results provide new insights into the assembly and mechanism of the tRNA ligase complex, shedding light on its functions in tRNA biogenesis and beyond.

## Introduction

Transfer RNA (tRNA) splicing is an essential and evolutionarily conserved process required for generating mature, functional tRNA molecules from primary transcripts ^1,2^. The human genome encodes 416 tRNA genes, 24 of which contain introns. This group includes five Arg (TCT), five Leu (CAA), one Ile (TCT), and thirteen Tyr (GTA) tRNA genes ^3,4^. In all cases, the intron is located one nucleotide 3′ to the anticodon. Correct intron removal and exon ligation are essential for proper folding into the characteristic L-shaped tertiary structure and for generating the correct anticodon loop, both of which are critical for translation. While tRNA introns are found in Archaea and most eukaryotes ^5,6^, the mechanisms for their removal and subsequent exon ligation vary among organisms. In human cells, the splicing process initiates with cleavage of the intron-containing pre-tRNAs by the tRNA splicing endonuclease (TSEN) complex, which generates cleaved tRNA halves bearing specific termini: a 2’,3’-cyclic phosphate and a 5’-hydroxyl group ^7^. Subsequently, these termini are ligated by the human tRNA ligase complex (tRNA-LC), a multi-subunit enzyme complex responsible for the second step of the splicing ^8^.

The human tRNA-LC comprises five subunits: the catalytic ligase subunit RTCB, and four auxiliary proteins: CGI-99, FAM98B, DDX1, and Ashwin (ASW) ^8,9^. RTCB catalyzes the ligation step by directly sealing the 2’,3’-cyclic phosphate and 5’-hydroxyl termini of cleaved tRNA molecules, a unique enzymatic mechanism conserved in Archea and Metazoa, and distinct from ligases found in plants and fungi ^10–12^. CGI-99 and FAM98B form a heterodimer essential for maintaining structural integrity of the complex ^13^. The DEAD-box RNA helicase DDX1 has been implicated in the regulation of tRNA-LC activity and may facilitate RNA remodeling during splicing, although its precise mechanistic role remains to be fully elucidated ^9^. Beyond its role in the tRNA-LC, DDX1 has been shown to participate in diverse cellular processes, including mRNA transport, DNA double-strand break repair, and miRNA processing, suggesting that it may integrate RNA metabolism with broader cellular regulatory networks ^14–16^. ASW, a vertebrate-specific subunit, has been proposed to stabilize the overall architecture of tRNA-LC but is dispensable for RTCB binding, suggesting a supportive rather than catalytic role in the complex ^13^. In addition, two regulatory factors, Archease and PYROXD1, are critical for tRNA-LC function ^17–19^. Archease is required to guanylylate a key catalytic histidine residue in the RTCB active site, priming the enzyme for the ligation reactions ^18^. PYROXD1, an NADPH-dependent oxidoreductase, binds RTCB in an NAD(P)H-dependent manner and protects its active site from oxidative inactivation ^17,20^. Importantly, Archease and PYROXD1 binding is mutually exclusive with substrate engagement, as both occupy the RTCB active site and block tRNA binding^17,19,20^.

Dysfunction of tRNA processing enzymes is increasingly recognized as a contributor to various human diseases, underscoring the need for detailed structural and mechanistic insights into this critical pathway. Mutations in PYROXD1 cause a muscular dystrophy-like phenotype ^21^, while mutations in TSEN subunits as well as the RNA kinase Clp1 lead to pontocerebellar hypoplasia subtypes ^21,22^. Furthermore, DDX1 is frequently upregulated in various cancers ^14^. Beyond its function in tRNA maturation, RTCB plays a critical role in the unfolded protein response (UPR) by facilitating the unconventional splicing of *XBP1* mRNA in conjunction with the UPR sensor IRE1, producing an alternatively spliced isoform that drives UPR signaling ^23–26^. DDX1 has further been linked to HIV replication and is reported to participate in multiple distinct protein complexes ^27,28^.

High-resolution structures of RTCB in complexes with Archease ^19^ and PYROXD1 ^20^ have been elucidated, revealing the molecular details of their activation and protection mechanisms. However, the structural organization of the tRNA-LC remains poorly understood, despite its fundamental importance in RNA biology and disease. Previous structural and biochemical studies have provided preliminary insights into the overall architecture and interactions within human tRNA-LC ^13,19,20^, yet critical details remain unresolved, in particular the molecular determinants underpinning complex assembly and the functional roles of auxiliary subunits in modulating complex stability and activity. Furthermore, the existence of multiple paralogs of the tRNA-LC subunit FAM98B raises intriguing questions about their potential functional specialization and redundancy within human cells.

In this study, we utilized cryo-electron microscopy (cryo-EM) complemented by targeted biochemical and genetic experiments to elucidate the high-resolution structure of the human tRNA-LC. Our findings define the core structural determinants required for complex assembly, characterize essential inter-subunit interfaces, and provide mechanistic explanations for previous functional data. Importantly, we identify novel compositionally distinct complexes involving paralogous FAM98 subunits, revealing previously unappreciated functional diversity of human RTCB ligase complexes. Together, these structural and mechanistic insights advance our understanding of human tRNA processing and set the foundation for investigating how alterations in tRNA-LC components may contribute to human diseases.

## Results

### Overall architecture of the five-subunit human tRNA-LC

To determine the three-dimensional structure of human tRNA-LC, we generated a truncated variant of the tRNA-LC complex by co-expression of all five subunits in baculovirus-infected insect cells. While FAM98B, CGI-99, and DDX1 were stably associated with RTCB, ASW was initially recovered in sub-stoichiometric amounts, necessitating a tandem-affinity purification approach using a Twin-Strep tag fused to the C-terminus of Ashwin. To facilitate structural analysis by cryo-EM, we reconstituted a minimal five-subunit tRNA-LC (**Fig. 1A, Fig.S 1**) comprising full-length RTCB, CGI-99 and ASW subunits together with a truncated FAM98B construct (comprising residues 1-300) that lacked an intrinsically disordered C-terminal region and a C-terminal fragment of DDX1 (residues Gly696-Phe740^DDX^^1^), which was previously shown to be sufficient for tRNA-LC assembly. The DDX1 RecA1, RecA2 and SPRY domains were omitted as their presence impaired particle alignment during preliminary cryo-EM analysis of full-length tRNA-LC. To overcome preferential orientation of tRNA-LC particles, we employed blotless vitrification and merged untilted data with an additional particle dataset acquired at high tilt angles, ultimately obtaining an atomic reconstruction at a nominal resolution of 3.3 Å (**Fig. 1B,C; Fig. S2,4**). While the CGI-99, FAM98B, DDX1 and Ashwin subunits were well resolved, the map lacked definition in the region corresponding to RTCB, likely due to residual particle orientation bias. To address this, we additionally determined a cryo-EM reconstruction of the minimal tRNA-LC complex described above bound to PYROXD1 in the presence of NADH, at a nominal resolution of 3.3 Å (**Fig. 1D,E; Fig. S3,4**). The resulting map had improved local resolution in the RTCB region, enabling us to build a composite atomic model for the entire tRNA-LC starting from available crystal structures and AlphaFold3 (AF3) structural models ^29^.

**Fig 1.**
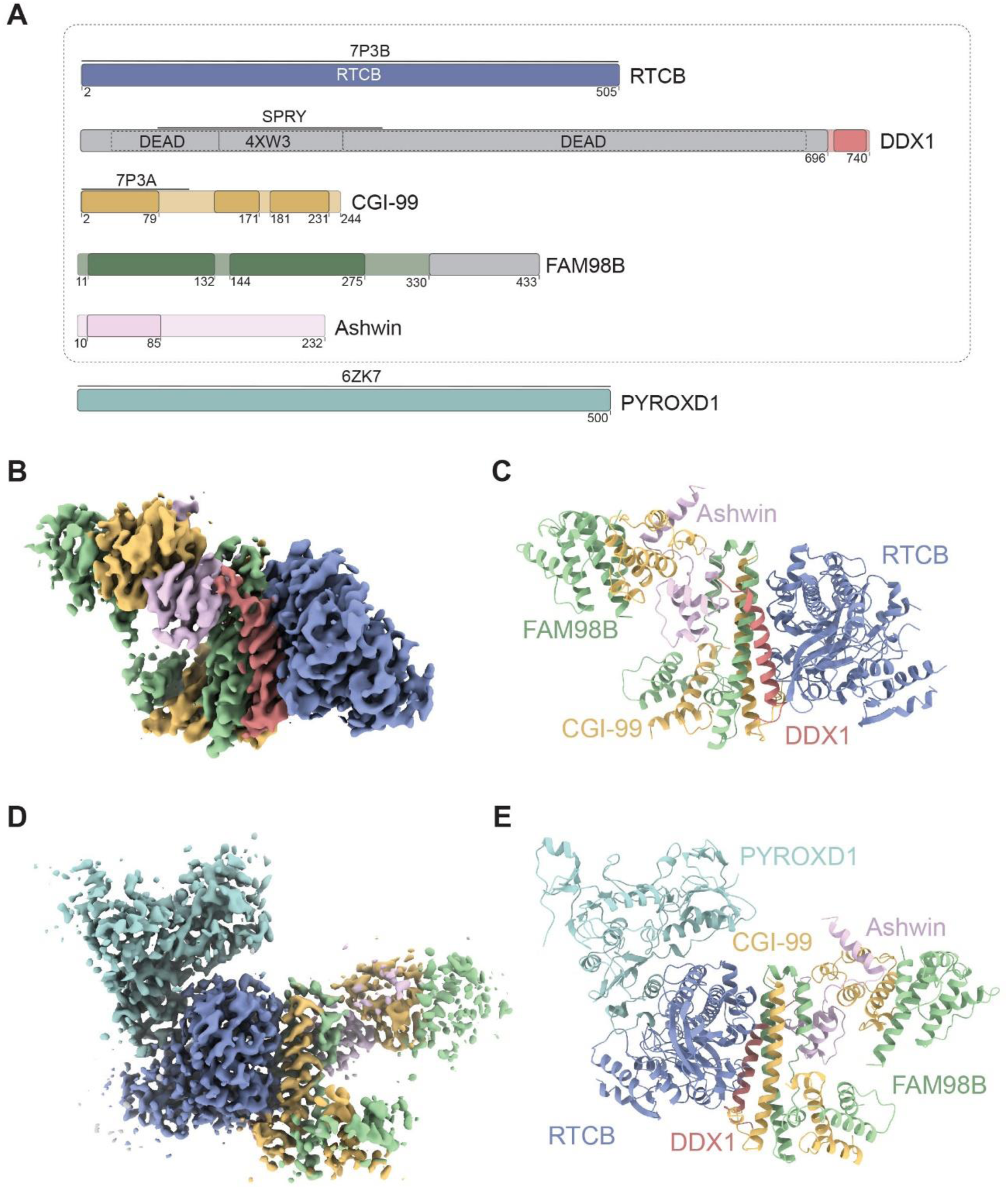
Structures of the human tRNA-LC. (**A**) Domain organization of tRNA-LC subunits and PYRODX1. Structured regions of the proteins are highlighted. Previously determined structures are indicated with their respective PDB codes. (**B**) cryo-EM reconstruction of the tRNA-LC. (**C**) Atomic model of tRNA-LC corresponding to the reconstruction in (B). (**D**) cryo-EM reconstruction of the tRNA-LC. (**E**) Atomic model of tRNA-LC corresponding to the reconstruction in (D).

The structure reveals that the minimal structural core of human tRNA-LC is formed by the globular ligase subunit RTCB and its interface with a four-helical bundle formed by conserved C-terminal regions of CGI-99, FAM98B and DDX1, in agreement with our prior biochemical analysis of human tRNA-LC. CGI-99 and FAM98B co-fold into an intricate heterodimeric subcomplex with two independent modules, hereafter termed the head and interface modules (**Fig. 2A**). The architecture of the FAM98B-CGI-99 heterodimer is consistent with prior data showing that the two proteins are capable of forming a stable heterodimer in the absence of the other tRNA-LC subunits. The head module comprises the N-terminal calponin homology (CH)-like domains (residues Phe2–Gly93^CGI-99^ and Thr11–Asn132^FAM98B^), while the interface module comprises the alpha-helical C-terminal regions of both subunits (residues Pro128–Ala231^CGI-99^ and Ser144–Arg275^FAM98B^) and includes structural elements mediating interactions with RTCB and DDX1. Together, the two modules adopt a pincer-like configuration to form the binding interface for the structured N-terminal domain of ASW (residues Cys10–Asn85^ASW^). The C-terminal region of ASW (residues Glu86-Pro232^ASW^) is disordered and not resolved in our model, in agreement with AF3 structural modeling. There are no contacts between Ashwin and RTCB, validating previous interaction data indicating that Ashwin is not required for the interaction of RTCB with the other tRNA-LC subunits.

**Fig 2.**
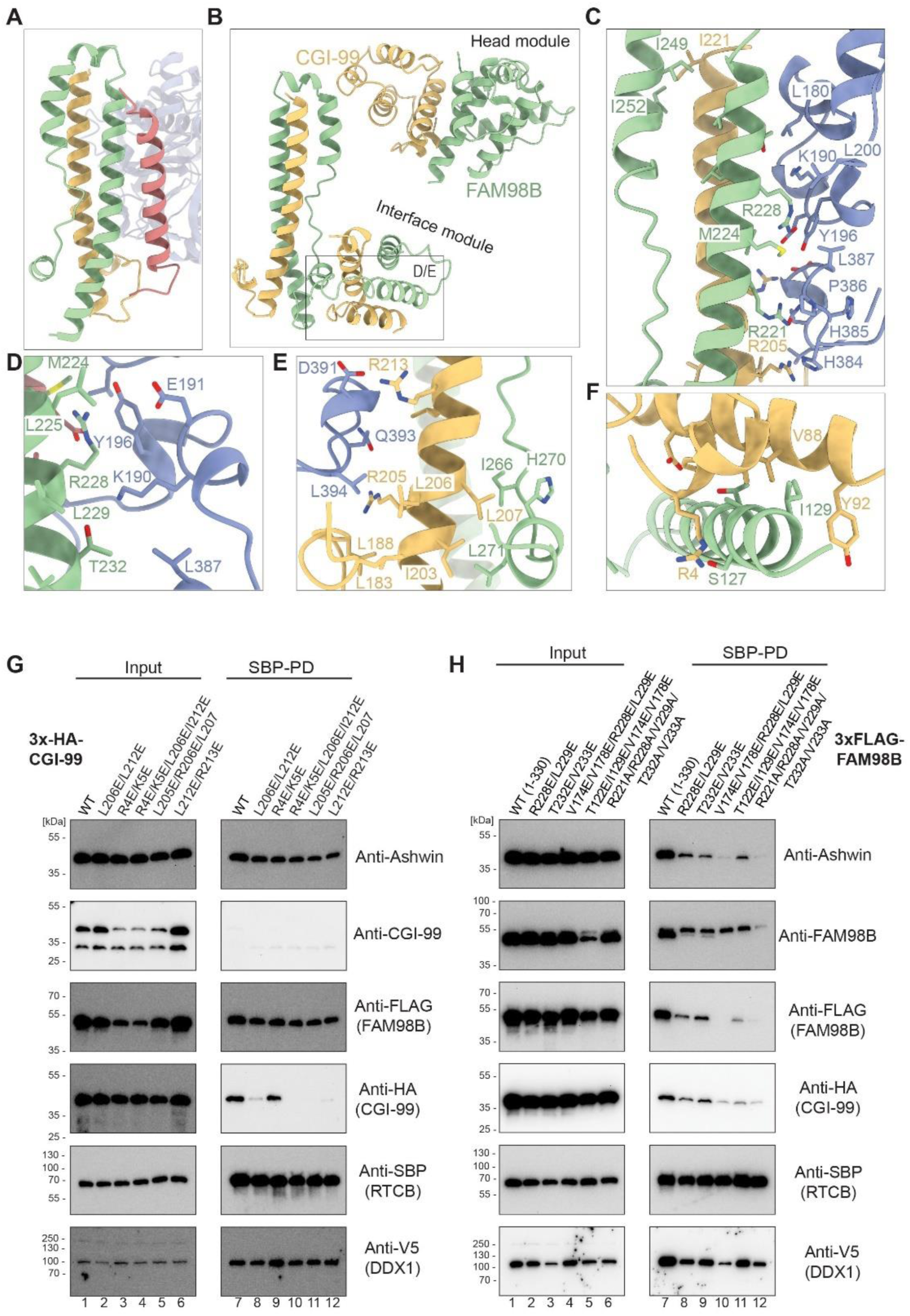
FAM98B and CGI-99 form an intricate heterodimeric subcomplex. (**A**) Cartoon representation of the FAM98B/CGI-99 dimer. Colors correspond to the coloring scheme in Fig. 1. The head and interface modules are indicated. (**B–F**) Close-up views showing key interface residues involved in the CGI-99–FAM98B interaction. (**G and H**) Streptavidin affinity co-precipitation of wild-type (WT) and mutant FAM98B or CGI-99 proteins co-expressed with the other tRNA-LC subunits in HEK293T cells. SBP-tagged RTCB was used as bait.

Notably, the ligase catalytic site is located on the opposite side of RTCB, away from the tRNA-LC core interface. No contacts are observed between PYROXD1 and the four non-catalytic subunits in the tRNA-LC-PYROXD1 complex, indicating that PYROXD1 is recruited to RTCB independently of CGI-99, FAM98B, DDX1 and Ashwin. Likewise, structural superposition of the tRNA-LC with the recently determined structure of the RTCB-Archease complex suggests that the non-catalytic tRNA-LC subunits do not contribute to Archease recruitment.

### FAM98B, CGI-99 and DDX1 interface with RTCB via a helical bundle

Our previous interaction studies revealed that the C-terminal regions of FAM98B, CGI-99 and DDX1 are required for the structural integrity of the human tRNA-LC. In the present structure, the respective C-terminal regions of the subunits (residues Asn200–Arg275^FAM98B^, Ala195–Ala231^CGI-99^ and Val705–Leu736^DDX^^1^) form a four-helix bundle, with FAM98B contributing two alpha-helices flanked by one helix each from CGI-99 and DDX1. All three subunits engage with RTCB via one helix, interacting with an extensive patch that includes residues His384-Leu387^RTCB^, Asp391^RTCB^, Gln393^RTCB^, Leu394^RTCB^, Leu180^RTCB^, Lys190^RTCB^, Glu191^RTCB^, Tyr196^RTCB^. Inter-helical interactions responsible for the structural integrity of the helical bundle are mediated by an intricate network or hydrophobic and salt-bridge interactions centered on invariant or highly conserved residues (**Fig. 2B–F**). For CGI-99, these include residues Arg205^CGI-99^, Leu206^CGI-99^, Leu207^CGI-99^, Leu212^CGI-99^ and Arg213^CGI-99^, involved in contacts with RTCB and/or CGI-99. For FAM98B, the interaction network includes residues Arg221^FAM98B^, Met224^FAM98B^, Leu225^FAM98B^, Arg228^FAM98B^, Thr232^FAM98B^ and Val233^FAM98B^.

To support our structural observations, we co-expressed structure-based mutant CGI-99 and FAM98B proteins together with the other tRNA-LC subunits in HEK293T cells and performed streptavidin co-precipitation assays using Streptavidin Binding Peptide (SPB)-tagged RTCB as bait. Due to the extensive nature of the inter-subunit interfaces, we tested combinations of mutations focused on strictly conserved residues. CGI-99 proteins containing specific mutations within the RTCB/FAM98B interaction interface were not co-precipitated with RTCB, confirming the importance of these residues for tRNA-LC assembly (**Fig. 2G**). In contrast, mutations of residues mediating FAM98B interactions within the head module of the FAM98B-CGI-99 heterodimer did not impact tRNA-LC assembly. For FAM98B, the introduction of double point mutations in the interface helix reduced co-precipitation efficiency (**Fig. 2H**). Complete loss of FAM98B interaction required a more extensive combination of mutations within the interface helix (R221A/R228A/V229A/T232A/V233A) or their combination with mutation of other residues meditating contacts with CGI-99 (V174E/V178E/R228E/L229E). Notably, the expression of FAM98B mutants perturbed the co-precipitation efficiency of CGI-99 and ASW. This observation can be explained by sequestration of Ashwin and CGI-99 by the mutant FAM98B proteins and is in agreement with prior observations that CGI-99, FAM98B and ASW form a stable trimeric complex in isolation ^13^. Together, these structural observations and biochemical results define key molecular determinants within CG-99 and FAM98B that are critical for tRNA-LC assembly and integrity.

### DDX1 is recruited to tRNA-LC via its C-terminal helix

The C-terminal region of DDX1 spanning residues Gly696-Phe740^DDX^^1^ was previously shown to be necessary and sufficient for assembly of a minimal human tRNA-LC. The tRNA-LC structure reveals that this region comprises a highly conserved single alpha-helix, hereafter termed the C-terminal helix (CTH), within the core interface four-helix bundle (**Fig. 3A,B**). A constellation of invariant residues within the CTH engage in interactions with FAM98B and/or RTCB (**Fig. 3B–D**), as well as CGI-99 (**Fig. 3E**). Leu715^DDX^^1^, Leu718^DDX1^ and Glu719^DDX1^ primarily interact with FAM98B. In turn, Gln723^DDX1^, Phe726^DDX1^ and L727^DDX1^ make hydrogen-bonding and hydrophobic contacts with RTCB, while L729^DDX1^ and L736^DDX1^ interact with CGI-99.

**Fig 3.**
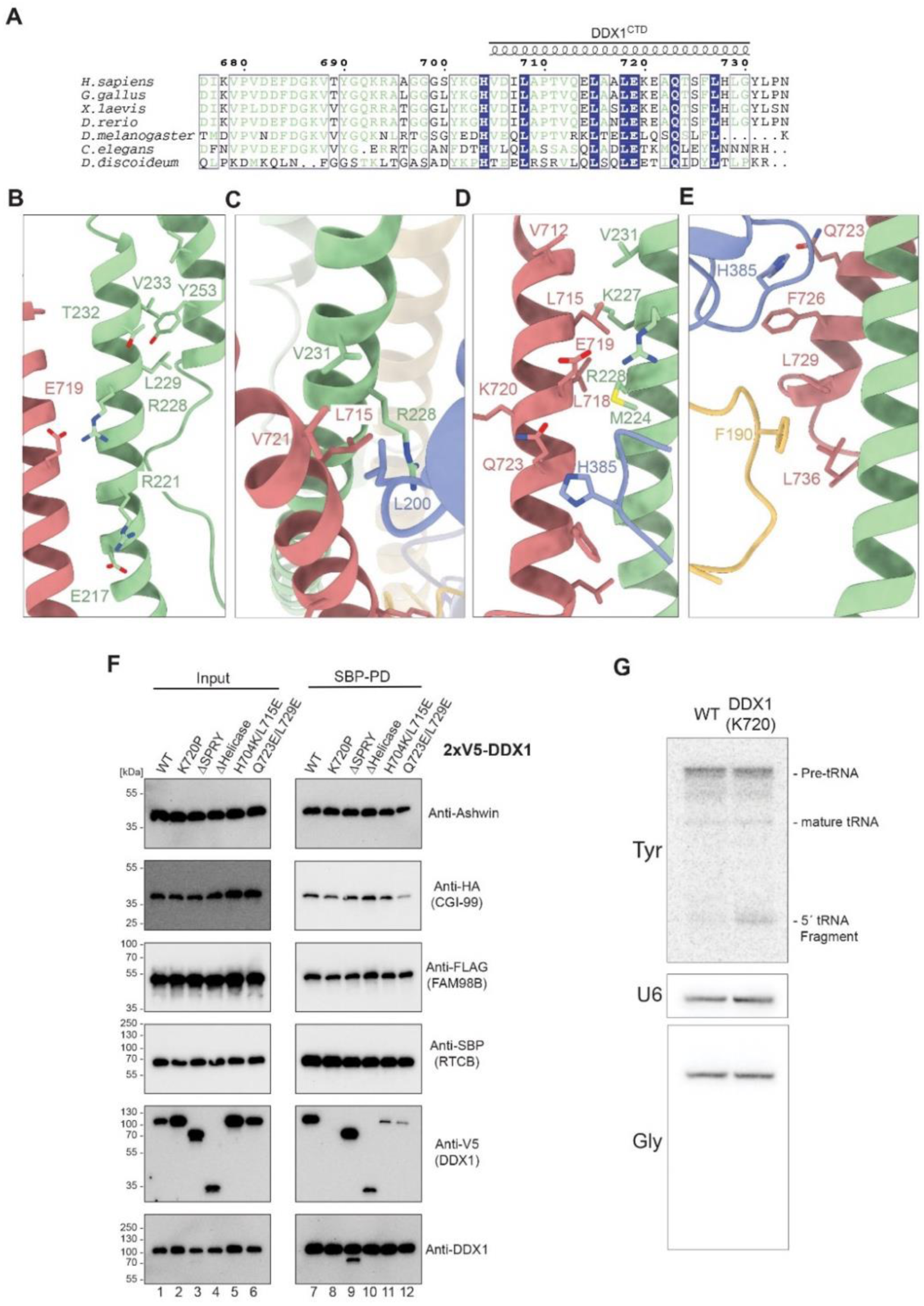
C-terminal helix recruits DDX1 to tRNA-LC. (**A**) Multiple sequence alignment of the C-terminal regions of DDX1 orthologs. The C-terminal helix, as observed in the tRNA-LC structure is indicated above the sequences. (**B-E**) Zoom-in view of the helical bundle formed by FAM98B, CGI-99 and DDX1. Subunit color corresponding to the coloring scheme in Fig. 1. (**F**) Streptavidin affinity co-precipitation of wild-type (WT) and mutant DDX1 proteins co-expressed with the other tRNA-LC subunits in HEK293T cells. SBP-tagged RTCB was used as bait. (**G**) Northern blot analysis of tRNA maturation in WT and homozygous K720P^DDX1^ mutant HEK293T cells. The presence of a 5’-fragment band in the tRNA^Tyr^ blot indicates a splicing defect. U6 RNA and tRNA^Gly^ serve as loading controls.

To validate the importance of the CTH for the assembly of DDX1 within the tRNA-LC, we tested truncated or point-mutant DDX1 constructs in the SBP-RTCB co-precipitation assay. Disruption of the CTH by a proline substitution of Lys720^DDX1^ resulted in complete loss of DDX1 co-precipitation, while deletion of either the helicase domains or the SPRY domain insertion had no impact on DDX1 recruitment (**Fig. 3F**). Furthermore, point mutations of strictly conserved residues resulted in substantial loss of DDX1 co-precipitation. Together, these results confirm the DDX1 CTH as the sole interaction motif responsible for the assembly of DDX1 within the tRNA-LC.

Previous studies have shown that DDX1 is required for tRNA biogenesis, as genetic knockout of DDX1 in the human U2OS cell line results in the accumulation of unspliced pre-tRNA intermediates ^30^. To test whether the function of DDX1 in pre-tRNA splicing is dependent on its association with RTCB within tRNA-LC, we generated a HEK293T cell line carrying a homozygous DDX1 K720P mutation, which selectively disrupts DDX1 assembly within the tRNA-LC and analyzed tRNA^Tyr^ processing using Northern blotting. This experiment revealed marked accumulation of unspliced tRNA fragments in the K720P mutant line, as compared to wild-type cells (**Fig. 3G**), indicating that the association of DDX1 within tRNA-LC is required for proper tRNA maturation.

### Ashwin bridges the head and interface modules of FAM98-CGI-99

ASW is wedged in between the head and interface modules of the FAM98B-CGI-99 heterodimer (**Fig. 4A**), extensively interfacing with the CH domain of CGI-99 in the head module (1118 Å^2^) and the interface helices of FAM98B (593 Å^2^). The structured part of ASW comprises a compact N-terminal helical domain (residues Cys10-Ile58^ASW^), which is followed by a structured linker (Pro59-Arg69^ASW^) that threads through a gap between the head and interface modules and a single-alpha helix terminating at Asn85^ASW^ (**Fig. 4B**). Key interaction residues include Ile58^ASW^, Pro59^ASW^, Pro61^ASW^, Arg63^ASW^, Trp70^ASW^ and Lys77^ASW^ (**Fig. 4C**). Validating the observed interaction mode using the SBP-RTCB co-precipitation, point mutations of Pro59^ASW^, Trp70^ASW^ or Lys77^ASW^ completely abolished ASW assembly within tRNA-LC, as did substitution of the Pro59-Arg69^ASW^ linker with a Gly-Ser dipeptide (**Fig. 4D**). These results confirm that ASW assembly in the tRNA-LC is solely dependent on interactions with FAM98B and CGI-99, while the integrity of tRNA-LC is not disrupted by the loss of ASW binding.

**Fig 4.**
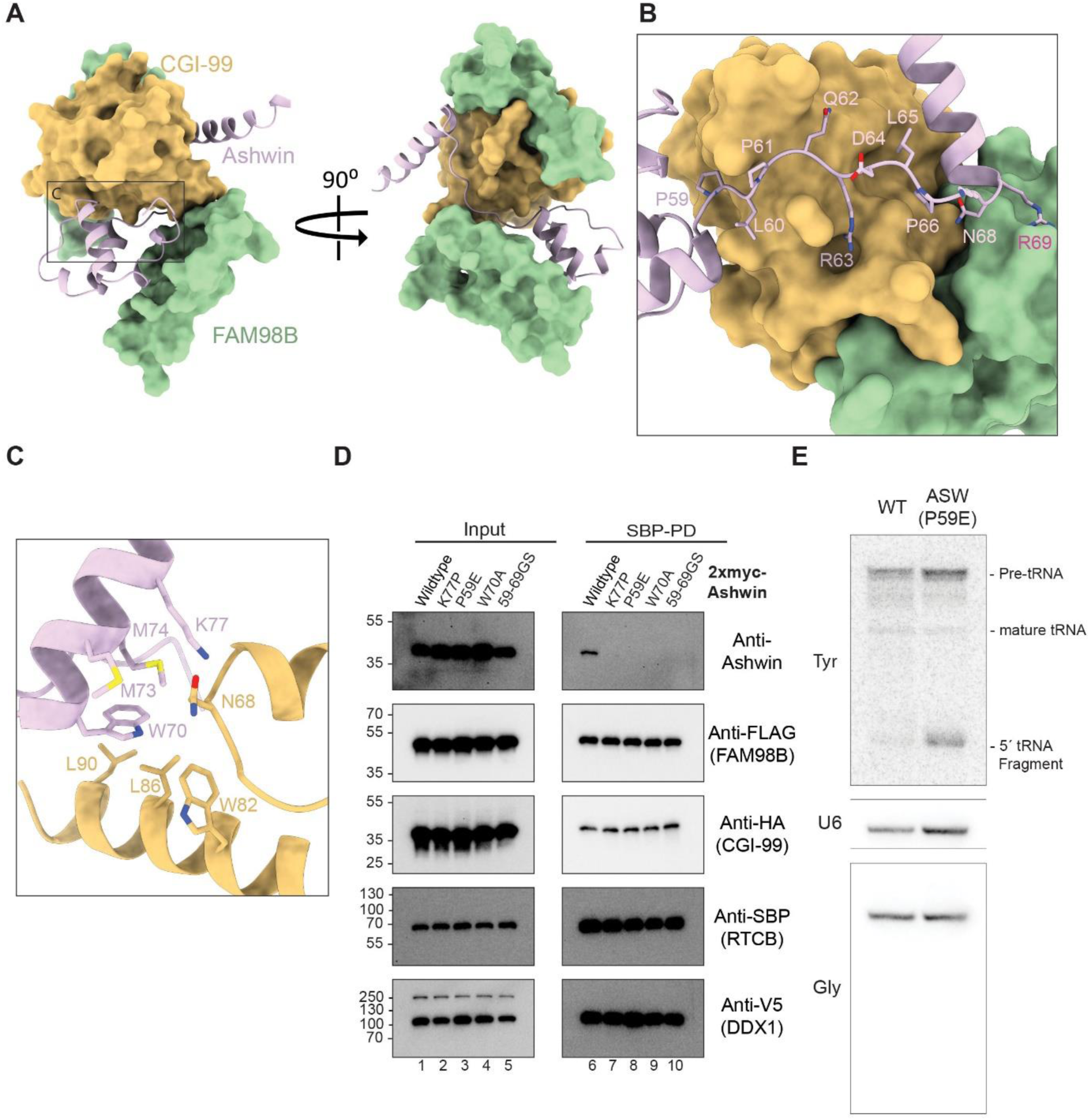
Ashwin bridges the head and interface modules of FAM98-CGI-99. (**A**) Overview of Ashwin clamped by the head and interface modules the FAM98B-CGI-99 subcomplex. (**B**) Close-up view of key residues at the interface of Ashwin and CGI-99. (**C**) Close-up view of the extended helix (residues 60-77^ASW^) of Ashwin contacting CGI-99. (**D**) Streptavidin affinity co-precipitation of wild-type (WT) and mutant Ashwin proteins co-expressed with the other tRNA-LC subunits in HEK293T cells. SBP-tagged RTCB was used as bait. (**E**) Northern blot analysis of tRNA maturation in WT and homozygous P59E^ASW^ mutant HEK293T cells. The presence of a 5’-fragment band in the tRNA^Tyr^ blot indicates a splicing defect. U6 RNA and tRNA^Gly^ serve as loading controls.

To test whether the incorporation of ASW in the tRNA-LC is required for pre-tRNA splicing, we generated a homozygous HEK293T cell line carrying the W59P^ASW^ mutation, designed to specifically disrupt its recruitment to the tRNA-LC. Northern blotting analysis revealed accumulation of unspliced tRNA^Tyr^ fragments. This result indicates that the assembly of ASW into tRNA-LC is required for pre-tRNA splicing, thus confirming the functional importance of the integrity of the pentameric tRNA-LC holocomplex for tRNA biogenesis.

### FAM98 paralogs assemble into compositionally and functionally distinct RTCB ligase complexes

Our structural analysis establishes FAM98B as a key structural component of human tRNA-LC. Mammalian genomes encode three paralogous FAM98 proteins: FAM98A, FAM98B and FAM98C. The high degree of amino acid sequence conservation within the predicted structured regions of FAM98A and FAM98C (54% between FAM98B and FAM98A, and 32% between FAM98B and FAM98C within residues 1-330) suggests they likewise interact with RTCB and the other tRNA-LC subunits. To test this hypothesis, we overexpressed 3xFLAG-tagged FAM98A/B/C proteins in HEK293T cells, performed anti-FLAG immunoprecipitation, followed by mass spectrometry analysis to identify interacting proteins. For all three FAM98 paralogs, tRNA-LC subunits RTCB, DDX1 and CGI-99 were among the most abundant co-purifying proteins (**Fig. 5A**). Notably, however, ASW did not co-purify with FAM98A and FAM98C. To validate these findings, we reconstituted FAM98A- and FAM98C-containing RTCB complexes by co-expression with RTCB, CGI-99 and DDX1 in insect cells and affinity purification (**Fig. 5B**). Together, these results indicate that FAM98A and FAM98C assemble in RTCB-containing, tRNA-LC-like complexes that lack ASW and are thus compositionally distinct from the canonical, FAM98B-containing tRNA-LC.

**Fig 5.**
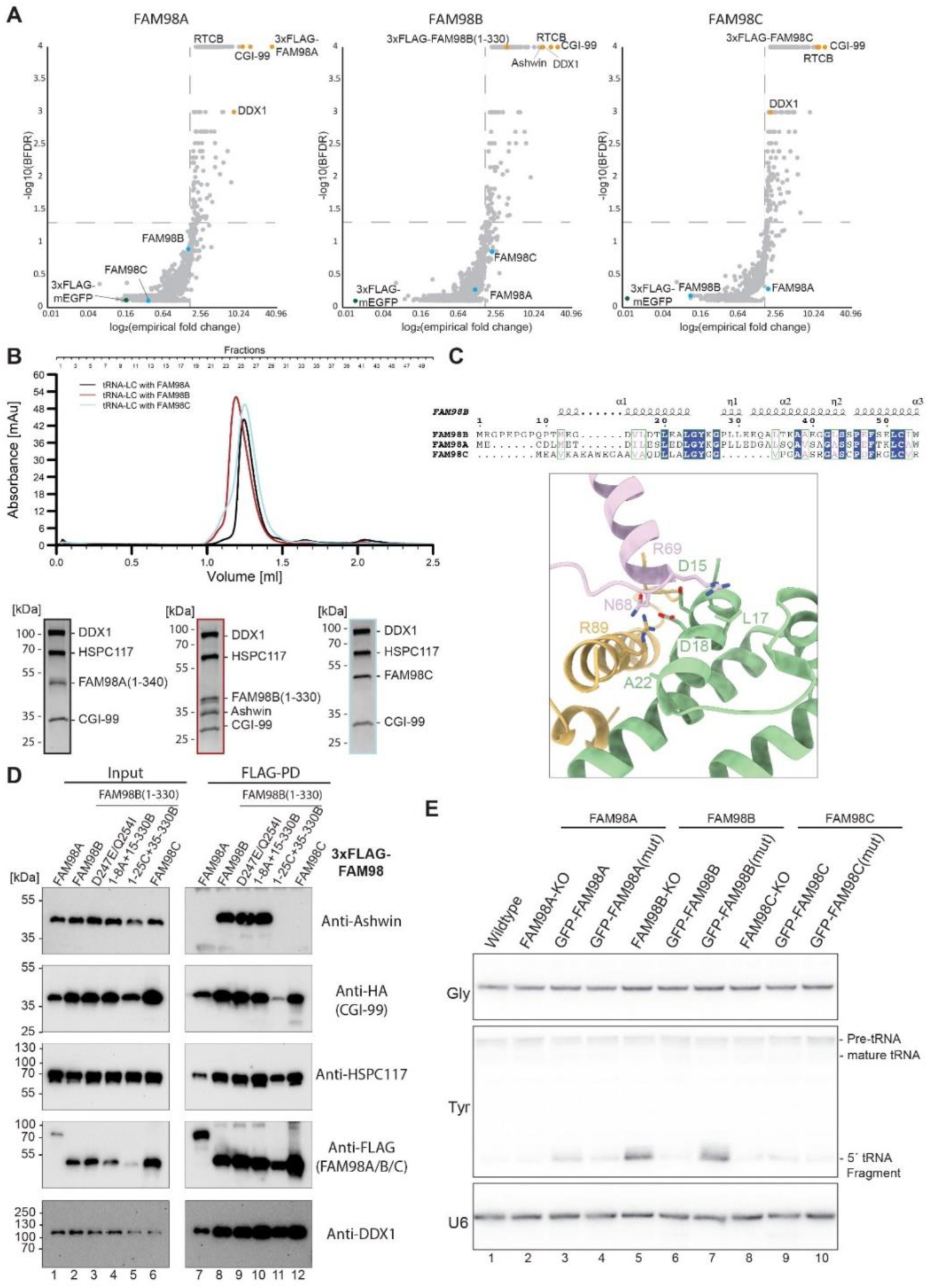
FAM98B is required for ASW assembly into tRNA-LC and tRNA processing. (**A**) Interaction proteomics of FAM98 paralogs, as based on affinity purification using 3xFLAG-tagged FAM98A/B/C proteins as bait versus mEGFP as negative control. (**B**) Size exclusion chromatography (top) and SDS-PAGE analysis (bottom) of recombinantly expressed RTCB complexes containing FAM98A/B/C subunits. (**C**) Multiple sequence alignment of the N-terminal region of FAM98A/B/C and closeup on the contacts mediated by this region with CGI-99 and Ashwin. (**D**) Streptavidin affinity co-precipitation of wild-type (WT) and mutant DDX1 proteins co-expressed with the other tRNA-LC subunits in HEK293T cells. SBP-tagged RTCB was used as bait. (**E**) Northern blot analysis of tRNA maturation in FAM98A/B/C CRISPR-knockout cell lines stably transfected with wild-type and mutant FAM98 expression constructs. The presence of a 5’-fragment band in the tRNA^Tyr^ blot indicates a splicing defect. U6 RNA and tRNA^Gly^ serve as loading controls.

Close inspection of the ASW-FAM98B interaction interface reveals that the FAM98B N-terminal helix mediates extensive contacts with ASW (**Fig. 5C**). The FAM98 paralogs differ principally in their N-termini (**Figure 5C**). FAM98C, in particular, contains a six-residue insertion within the first α-helix (**Fig. 5C**) as well as a deletion spanning residues Gly27–Leu35^FAM98B^. To define the structural determinants underpinning the specific association of ASW with FAM98B-containing tRNA-LC, we performed co-precipitation experiments using SBP-tagged FAM98A-C protein constructs in HEK293T cells. Validating our proteomic analysis, ASW efficiently co-precipitated with FAM98B, along with other tRNA-LC components, while neither FAM98A nor FAM98C supported ASW association (**Fig. 5D**). Replacement of the N-terminal residues Met1-Leu35 in FAM98B with the corresponding sequence from FAM98C (Met1-Gly25) resulted in loss of ASW co-precipitation (**Fig. 5D**), confirming the importance of the FAM98B N-terminal helix for ASW recruitment. Mimicking FAM98A, neither the substitution of FAM98B residues 1-8 with the corresponding amino acid sequence from FAM98A, nor the double mutation D247E/Q254I were sufficient to disrupt ASW co-precipitation (**Fig. 5D**). This implies that that the inability of FAM98A to support ASW recruitment likely results from a cumulative effect of multiple amino-acid substitutions. Taken together, our data indicates despite their substantial sequence conservation, the differences between the FAM98 paralogs result in subtle structural perturbations that have a profound effect on their interactions with ASW.

The canonical, FAM98B-containing tRNA-LC has a well-established function in pre-tRNA splicing. To shed light on the cellular functions of FAM98A- and FAM98C-containing RTCB complexes, we sought to test whether FAM98A or FAM98C complexes are involved in tRNA biogenesis by analyzing pre-tRNA^Tyr^ splicing in cells in which the assembly of FAM98A/B/C with RTCB is impaired. To this end, we generated stably transfected cell lines in which endogenous *FAM98A/B/C* gene expression was disrupted by CRISPR-genome editing and simultaneously complemented with expression of either wild-type FAM98 proteins or structure-based mutants incapable of RTCB association (**Fig. S5**). Disruption of the *FAM98B* gene resulted in the accumulation of unligated tRNA^Tyr^ fragments, indicating that FAM98B-containing tRNA-LC complex is essential for tRNA maturation. The misprocessing phenotype was rescued by overexpression of wild-type FAM98B not by expression of an association-deficient FAM98B mutant (R221A/R228A/V229A/T232A/V233A), providing further validation for the structural role of FAM98B in tRNA-LC assembly (**Figure 5E**). Disruption of endogenous *FAM98A* or *FAM98C* genes had no effect on tRNA^Tyr^ processing, suggesting that these paralogs are dispensable for tRNA ligation. Expression of WT FAM98A resulted in slight accumulation of misprocessed tRNA fragments whereas rescue with association-deficient FAM98A mutant had no effect on tRNA maturation, possibly due to competitive sequestration of RTCB by FAM98A. For *FAM98C* knock-out cells, rescue with WT FAM98C or association-deficient FAM98C mutant had an effect on tRNA maturation. Together, these results indicate that the integrity of the FAM98B tRNA-LC complex is essential for pre-tRNA processing in human cells, while FAM98A and FAM98C-complexes are not required for tRNA splicing, implying that they may have cellular functions distinct from tRNA biogenesis.

## Discussion

The human tRNA ligase complex is an essential RNA processing factor responsible for tRNA biogenesis and other cellular processes such as non-canonical *XBP1* mRNA splicing in the unfolded protein response, and RNA repair ^23,25,31^. In recent years, substantial progress has been made in understanding mechanistic aspects of its ligase subunit RTCB ^13^, including the catalytic priming by the coactivator Archease ^19^ and the protection from oxidative inactivation by PYROXD1 ^32^. However, structural details underpinning the assembly of RTCB within the human tRNA-LC have been elusive and hitherto limited to biochemical analyses based on cross-linking/mass-spectrometry and affinity co-precipitation experiments ^13^.

In our study, we present high-resolution cryo-EM reconstructions of the human tRNA-LC, complementing a recently determined structure of zebrafish tRNA-LC ^33^, and support our findings by interaction and functional *in cellulo* assays. The structure reveals an intricate architecture involving the cooperative assembly of the non-catalytic subunits FAM98B, CGI-99 and DDX1 into a helical bundle that interfaces with RTCB, explaining their co-dependency for tRNA-LC assembly ^13^. The structure further shows that the assembly of Ashwin into tRNA-LC depends on the co-folding of FAM98B and CGI-99 to generate a composite binding interface that does not involve RTCB, extending previous findings that Ashwin is not required for the assembly of the other tRNA-LC subunits ^13^. Notably, the ligase catalytic site on RTCB is spatially isolated from the subunit interfaces, underscoring the structural segregation of the catalytic activity from complex assembly. This implies that tRNA-LC’s non-catalytic subunits may not directly regulate RTCB catalytic activity, but rather modulate its cellular localization, stability, or recruitment of substrates.

In particular, DDX1 is required for tRNA biogenesis and has previously been shown to enhance Archease-dependent guanylylation of RTCB and its catalytic turnover in an ATP-dependent manner ^9,30^. Our structure reveals that the C-terminal helix of DDX1 is necessary and sufficient to mediate recruitment to the tRNA-LC, suggesting that DDX1 is flexibly tethered to the ligase. The flexibility likely serves to provide the helicase and SPRY domains of DDX1 with a dynamic radius of action, possibly to support Archease activity, as well as RNA substrate recruitment or release. We show that disrupting the recruitment of DDX1 to the tRNA-LC by single point mutations in the C-terminal helix leads to an accumulation of misprocessed tRNA fragments, confirming that tRNA biogenesis is dependent on the physical association of RTCB and DDX1. Notably, while DDX1 is required pre-tRNA processing, it appears to be dispensable for *XBP1* mRNA splicing. In light of our findings, this implies that the physical integrity of tRNA-LC is required for the nuclear function of RTCB in tRNA biogenesis but is not critical for its cytoplasmic role in the unfolded protein response. Moreover, although DDX1 is an integral tRNA-LC component, it has been implicated in a number of other processes ^15,16,28^. It is possible that the conformational flexibility of DDX1 enables its RTCB-independent functions while being assembled in the tRNA-LC. However, the precise functional roles of DDX1 within and outside of the tRNA-LC still remains enigmatic and warrants further investigation.

A key finding of our study is that the FAM98B paralogs, FAM98A and FAM98C, are also capable of assembling in RTCB complexes together with CGI-99 and DDX1. Although it might be expected that the FAM98 paralogs are functionally redundant, our proteomic analysis, supported by co-precipitation experiments, reveal that they are in fact compositionally distinct. While FAM98B supports the assembly of a heteropentameric tRNA-LC including Ashwin, the FAM98A/FAM98C RTCB complexes lack Ashwin. The compositional variability translates into functional divergence as well, with only the FAM98B complex being required for tRNA splicing. This points to a critical role of FAM98B and, by extension, ASW in the function of tRNA-LC in the context of tRNA biogenesis ^34^. As Ashwin has been predicted to harbor nuclear localization signals ^35^, our findings suggest that ASW enables the nuclear import of tRNA-LC, explaining the observed requirement of FAM98B for nuclear tRNA processing. Indeed, the complementary work of Leitner et al. identifies Ashwin as a nuclear import factor for the tRNA-LC. Taken together, these findings thus hint at the existence of multiple RTCB complexes, with a nuclear FAM98B complex involved in tRNA splicing and (cytoplasmic) FAM98A/FAM98C complexes that likely play alternative roles in RNA metabolism, possibly in UPR or RNA repair (**Figure 6**) ^36^.

**Fig 6.**
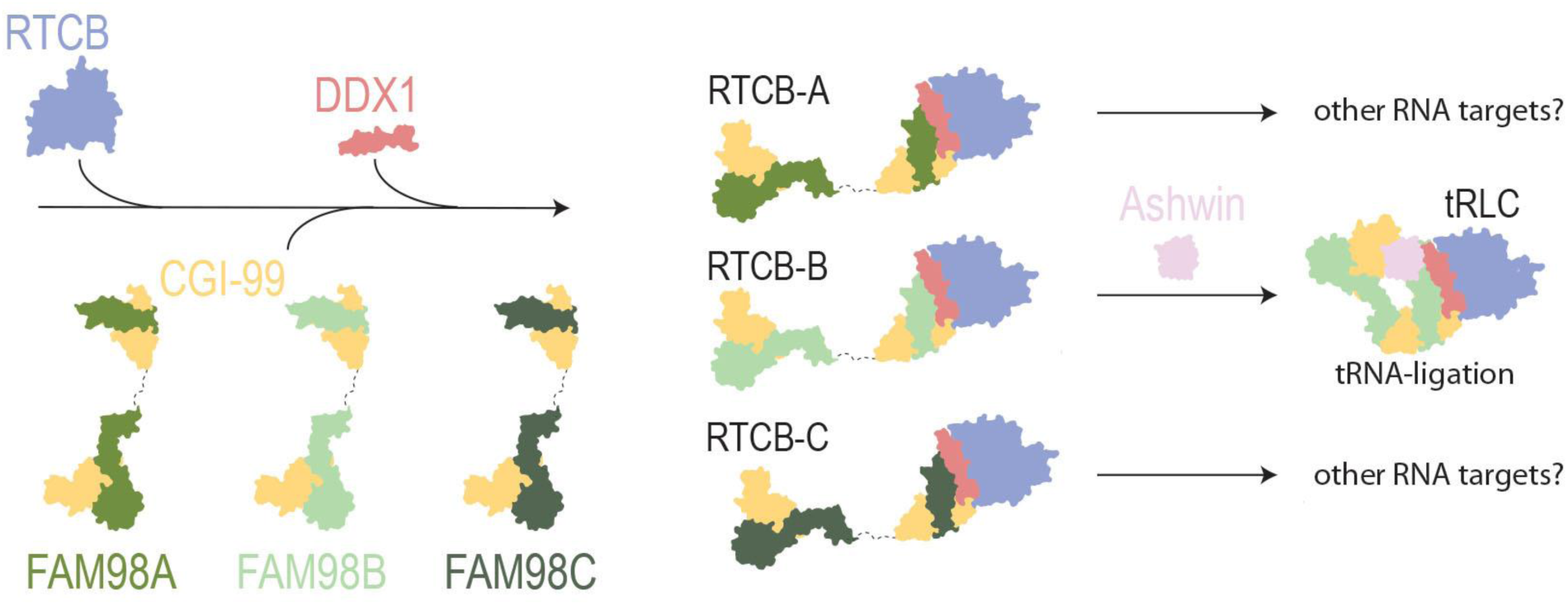
Compositional and functional diversity of human RTCB complexes. FAM98 paralogs underpin the assembly of compositionally and functionally distinct RTCB complexes. The FAM98B-containing RTCB complex includes ASW and is involved in nuclear tRNA processing. FAM98A-and FAM98C-containing complexes lack Ashwin and likely play other functional roles.

In conclusion, the presented work elucidates critical molecular determinants guiding the assembly, structural integrity, and compositional variability of human RTCB complexes. These findings provide a framework for future studies aimed at dissecting the regulatory mechanisms controlling tRNA splicing and broader functions of RNA ligase complexes in RNA metabolism beyond tRNA biogenesis.

## Acknowledgements

We are grateful to Piotr Szwedziak (University of Zurich Center for Microscopy and Image Analysis) and Emiko Uchikawa (Dubochet Center for Imaging, Lausanne) for technical support during cryo-EM sample preparation and data acquisition. We thank Judith Notbohm and Tina Perica for assistance with genome editing, and Stefano Pascarelli and Pedro Beltrao for phylogenetic analysis of FAM98 paralogs. We further thank all members of the Jinek lab for constructive feedback throughout the project. MMP was funded by the Swiss National Science Foundation (project no. TMPFP3_210571). MJ acknowledges funding by the Howard Hughes Medical Institute (International Research Scholar Award), the Vallee Foundation and the Swiss National Competence Center for Research “RNA & Disease”. Proteomic analysis was performed at the Functional Genomics Center Zurich (FGCZ) of the University of Zurich and ETH Zurich.

## Methods

### Cloning and expression of human tRNA-LC

Subunits of the tRNA-LC were cloned into modified and pre-tagged pLIB vectors ^37^. Ashwin was modified with a N-terminal 2xStrepII tag, RTCB with a N-terminal 8xHis tag and DDX1(696-740) was cloned together with C-terminal P2A-T2A-mEGPF tag. All subunits were assembled into a multi gene construct as described previously and used for recombinant expression in SF9 insect cells ^38^. Briefly, assembled multi gene cassettes were transformed into SF9 cells seeded in 6-well plates using TransIT®-Insect Transfection Reagent (Mirus, Cat. MIR 2304). 48 h after transfection the supernatant was removed, added to a 50 ml SF9 pre-culture and the cells incubated for further 48 h. Then the supernatant was removed again, adherent cells were mechanically detached from the 6-well plate and added to previous 50 ml SF9 pre-culture. 24 h later the pre culture was used to inoculate 750 ml expression cultures or multiple thereof at 1 mio. cells/ ml. The expression culture was incubated for 72 h at 27 °C. Infected insect cells were harvested by centrifugation (Heraeus, SLA4.1000, 500xg and 4 °C for 20 min), resuspended in lysis buffer (500 mM KCl, 20 mM HEPES-KOH, 30 mM Imidazole, pH 7.8), transferred to a 50 ml reaction tube and spun down again (Eppendorf, Rotor, 900 xg and 4 °C for 20 min). The supernatant was discarded and the pellet flash frozen in liquid nitrogen. Pellets were stored at -80 °C until further use.

### tRNA-LC purification

SF9 pellets with expressed tRNA-LC were resuspended in 5 volumes Lysis Buffer with 0.5 mM TCEP. Resuspended cells were lysed by sonication (Sonoplus, VS70 probe, 20 % amplitude, 2 min, 5 sec on/off cycle – repeated three times) and insoluble fractions were pelted by centrifugation (Hereaus, Rotor, 20,000 xg, at 4°C for 45 min). The clarified lysate was added to 0.1 volumes equilibrated Ni-NTA resin and incubated on a rolling table at 4 °C for 1 h. Subsequently, the resin-lysate slurry was centrifuged, for 1 min at 300 xg (Eppendorf), the supernatant was discarded, the resin resuspended in lysis buffer and added onto a gravity flow column. The resin was washed with multiple volumes Lysis Buffer. Next, bound tRNA-LC was eluted in multiple steps with 150 mM KCl, 20 mM HEPES-KOH, 250 mM Imidazole, pH 7.8. All elutions were pooled, added to 0.1 volumes equilibrated Streptactin resin (IBA) and incubated on a rolling table at 4 °C for 1 h. Subsequently the elution-resin slurry was centrifuged, resuspended in Wash Buffer 2 (150 mM KCl, 20 mM HEPES-KH, pH 7.8) supplemented with 0.5 mM TCEP and added to a gravity flow column. The resin was washed in several column volumes with Wash Buffer 2. Finally, the tRNA-LC was eluted in 150 mM KCl, 20 mM HEPES-KOH, 20 mM desthiobiotin, pH 7.8. All elutions were pooled, concentrated, aliquoted in 20 µl fractions and flash frozen in liquid nitrogen until further use. All steps of the purification were tracked by SDS-PAGE for quality assessment.

### Co-expression of tRNA-LC subunits in HEK293T cells

Full-length subunits of the tRNA-LC or truncations/ point mutants thereof were expressed using pre-tagged pCMV mammalian expression vectors. HEK293T cells were grown at 37 °C with 5% CO2 for all steps. Confluent HEK293T cells were trypsinized, seeded into 6-well plates (500,00 cells/ well) with 3 ml DMEM media supplemented with 10 % FBS, and incubated over night for recovery. The next day, 0.5 µg of each plasmid were mixed in 100 ul OptiMEM per well. In addition, 7.5 ug PEImax were mixed with 100 µl OptiMEM per well. PEImax and Plasmids were incubated for 5 min at room temperature and subsequently mixed. The PEImax – Plasmid mix was incubated for 30 min at room temperature and added dropwise onto each well. Subsequently, the cells were incubated for 72 h. Finally, cells were harvested by aspirating the media, addition of 1 ml Wash Buffer (150 mM KCl, 20 mM HEPES-KOH, pH 7.8) and detaching the cells by pipetting. Detached cells were transferred to 1.5 ml reaction tube, spun down (Eppendorf 5425 R, 900 xg for 5 min at 4 °C) and the supernatant was discarded. The cells were washed once as before and flash frozen in liquid nitrogen. Cell pellets were stored at -80 °C until further use.

### Streptavidin affinity co-purification assays

Harvested cells were resuspended in 500 µl Lysis Buffer 2 (150 mM KCl, 20 mM HEPES-KOH pH 7.8, 0.1% Tween20), lysed by sonication (MS72, 10 % Amplitude, 10 s) and centrifuged (30 min). 50 µl clarified lysate was mixed with 50 µl SDS-sample buffer and the remaining lysate was added onto 30 µl equilibrated Streptavidin resin. The resin-lysate mix was incubated at 4 °C on a turning wheel for 1 h. Subsequently, the resin was spun down at 500 xg and 4°C for 5 min, the supernatant removed and the resin washed 3 times as before with 180 µl Lysis Buffer 2. Finally, the resin was resuspended in 50 µl Lysis Buffer 2 and 50 µl SDS-sample buffer. Samples were stored at -20 °C until further use.

### Immunoblotting

Input and immunoprecipitation fractions were loaded onto gradient polyacrylamide gels (Biorad, AnyKd,) and separated at 300 V for 18 min. Proteins were transferred using preassembled transfer packs (Biorad, #1704156) with a Trans-Blot Turbo transfer system (Biorad). After transfer membranes were incubated in 5% Milk-TBST for 1 h on a rocking table. The milk-TBST solution was discarded, replaced with a milk-TBST antibody solution and again incubated for 1 h on a rocking table. Subsequently, the membrane was washed several times with TBST. Where appropriate, a secondary antibody in milk-TBST was added, again incubated for 1 h on a rocking temple and washed again with TBST. Finally, the immune blots were developed with ECL solution (Thermo Fisher) and imaged using a gel imager (Biorad).

### Sample vitrification for cryo-EM analysis

UltrAUfoil 300 1.2/1.3 or Quantifoil AU 300 1.2/1.3 grids were glow discharged for 30 s at 15 mA at 28 mbar. Purified protein was diluted to 0.3 mg/ ml 1.5 – 2.5 µl were applied to one side of the grid. Excess liquid was removed by blotting using Thermo Fisher Vitrobot Mark IV, with blot force 10 and 2 s blotting time. Subsequently the grid was plunge frozen in liquid ethane. Vitrified grids were stored in liquid nitrogen.

Alternatively purified tRNA-LC was adjusted to 18 mg/ ml with 1 mM FC8 and vitrified using self-wicking copper grids using a Chameleon instrument (SPT-Labtech) at the DCI Lausanne. The vitrification time was set to 501 ms.

### Cryo-EM Data collection and processing – tRNA-LC

Vitrified samples were imaged using a FEI Titan Krios G3i equipped with a Gatan Quantum Energy Filter and a Gatan K3 direct electron detector (4k x 4k) at the Center for Microscopy and Image Analysis at University of Zurich (UZH) and on a FEI Titan Krios equipped with a Gatan Quantum Energy Filter and a Falcon4i electron detector at the Dubochet Center for Imaging (DCI), Lausanne.

<u>Dataset #1 (UZH):</u> 7,307 movies were collected at 20 degree tilt angle with a total dose of 76 e/Å^2^ over 1.26 s and 60 frames. A defocus series of -0.8 nm to -2.4 nm was applied. The nominal magnification was set to 120,000. Data collection was performed in super-resolution mode with images recorded 2-fold binned resulting in a pixel size of 0.65 Å. Processing was done in CryoSPARC v4. Recorded movies were subjected to patch motion correction followed by patch CTF estimation ^39^. Particles picking was done using cryoSPARC internal tools yielding 8,620,162 identified particles which were extracted with 4-fold binning. After several rounds of 2D classification the remaining 691,104 particles were re-extracted at full pixel size after removing duplicates. After further 2D classification, the best 186,144 particles were selected and merged with dataset 02 for further processing.

<u>Dataset #2 (DCI):</u> 18,682 movies were collected with a total dose of 50 e/Å^2^ over 1.76 s and 50 frames. Dataset 02 was recorded on a self-wicking copper grid vitrified using the Chameleon system. A defocus series of -0.6 nm to -1.8 nm was applied. The nominal magnification was set to 120,000 with a pixel size of 0.65 Å. Recorded movies were subjected to patch motion correction followed by patch CTF estimation ^39^. Particles picking was performed using cryoSPARC internal tools yielding 4,303,118 identified particles which were extracted with 4-fold binning. After several rounds of 2D classification the remaining 190,994 particles were re-extracted at full pixel size after removing duplicates. After further 2D classification the best 137,765 particle were merged with the particles from Dataset 01. Pooled particles were used to generate an Ab-Initio model. The best Ab-Initio model was used to perform a Non-Uniform refinement which yielded a model of 3.25 Å from 291,506 particles. The map was further improved using DeepEMhancer ^40^. <u>Dataset #3 (UZH):</u> 9,536 movies were collected with a total dose of 60 e/Å^2^ over 1.26 s and 47 frames. A defocus series of -0.8 nm to -2.4 nm was applied. The nominal magnification was set to 120,000. Data collection was performed in super-resolution mode with images recorded 2-fold binned resulting in a pixel size of 0.65 Å. Processing was done in CryoSPARC v4. Recorded movies were subjected to patch motion correction followed by patch CTF estimation ^39^. Particles picking was done using cryoSPARC internal tools yielding 12,082,386 identified particles which were extracted with 4-fold binning. Several rounds of 2D classification identified a subset of assembled PYROXD1-tRNA-LC complexes consisting of 64,915 particles. Subsequent Ab-Initio model with 2 models and subsequent Non-Uniform refinement yielded two identical models which were merged and refined again. The resulting model was used for generation of templates for template picking which identified 5,912,851 particles. All particles were extracted with full pixel size and the best 20,131 particles were identified by 2D classification. The particles were merged Dataset 04 for further processing.

<u>Dataset #4 (UZH):</u> 10,153 movies were collected with a total dose of 62 e/Å^2^ over 1.26 s and 36 frames. A defocus series of -0.8 nm to -2.4 nm was applied. The nominal magnification was set to 120,000. Data collection was performed in super-resolution mode with images recorded 2-fold binned resulting in a pixel size of 0.65 Å. Processing was done in CryoSPARC v4. Recorded movies were subjected to patch motion correction followed by patch CTF estimation ^39^. Particles picking was done using cryoSPARC internal tools yielding 15,476,545 identified particles which were extracted with 4-fold binning. Several rounds of 2D classification identified 1,042,623 particles. After merging with particles from Dataset 03 Non-Uniform refinement was performed and followed by several rounds of 2D classification reducing the particle number to 129,417 which were merged with particles from dataset 05 and further 2D classified. Particles resembling the tRNA-LC bound to PYROXD1 were selected and used for homogenous refinement. The resulting 3D model was subjected to Heterogenous refinement resulting in a model of 269,780 particles. After global and local CTF refinement the particles were again subjected to Non-Uniform refinement yielding a final model at 3.3 Å.

<u>Dataset #5 (UZH):</u> 9,629 movies were collected with a total dose of 60 e/Å^2^ over 1.26 s and 47 frames. A defocus series of -0.8 nm to -2.4 nm was applied. The nominal magnification was set to 120,000. Data collection was performed in super-resolution mode with images recorded 2-fold binned resulting in a pixel size of 0.65 Å. Processing was done in CryoSPARC v4. Recorded movies were subjected to patch motion correction followed by patch CTF estimation ^39^. Particles picking was done using cryoSPARC internal tools yielding 12,866,744 identified particles which were extracted with 4-fold binning. After several rounds of 2D classification, the remaining 1,638,788 particles were subjected to Ab-Initio modelling. Models representing tRNA-LC related assemblies were selected and subjected to a further round of 2D classification. After removing duplicates and performing further 2D classifications, particles which resembled tRNA-LC bound to PYROXD1 were selected and merged with dataset 04.

### Proteomic analysis of FAM98 interactors

Modified pCMV vectors encoding N-terminally 3×FLAG-tagged FAM98A/B/C and 3×FLAG-mEGFP were expressed in HEK293T cells, immunoprecipitated using anti-FLAG resin, and analyzed by mass spectrometry. All samples were prepared in triplicate. Confluent HEK293T cells were trypsinized and seeded into 6-well plates at a density of 500,000 cells per well in 3 ml DMEM supplemented with 10% FBS, followed by overnight incubation to allow recovery. Subsequently, 2.5 µg of each plasmid were mixed with 100 µl OptiMEM per well. Separately, 7.5 µg PEImax were diluted in 100 µl OptiMEM per well. The PEImax and plasmid solutions were incubated separately for 5 minutes at room temperature, then combined and incubated for an additional 30 minutes. The resulting transfection mix was added dropwise to each well. Cells were then incubated for 72 hours (37 °C with 5% CO2). For harvesting, culture medium was aspirated, and 1 ml of Wash buffer (150 mM KCl, 20 mM HEPES-KOH, pH 7.8) was added per well. Cells were detached by pipetting and transferred to 1.5 ml reaction tubes. Samples were centrifuged (Eppendorf 5425 R, 900 × g, 5 min, 4 °C), the supernatant discarded, and the cells washed once more with wash buffer. Cell pellets were flash-frozen in liquid nitrogen and stored at –80 °C until further use. Frozen cell pellets were thawed and resuspended in 500 µl Lysis buffer 2 (150 mM KCl, 20 mM HEPES-KOH, pH 7.8, 0.1% Tween-20), lysed by sonication (MS72, 10% amplitude, 10 s), and clarified by centrifugation (30 min). The resulting supernatant was incubated with 30 µl anti-FLAG resin (ANTI-FLAG® M2 Affinity Gel, Sigma) on a rotating wheel for 1 hour at 4 °C. After incubation, the resin-lysate mixture was centrifuged (Eppendorf 5425 R, 900 × g, 5 min, 4 °C), the supernatant removed, and the resin washed three times with 200 µl Wash buffer. Finally, the resin was resuspended in 50 µl Wash buffer, and the samples were submitted for mass spectrometry analysis at the Functional Genomics Center Zurich.

Captured proteins were digested on beads using 500 ng of sequencing-grade modified Trypsin (Promega, V5111) including cysteine reduction and alkylation by Tris(2-carboxyethyl)phosphine hydrochloride (TCEP) and 2-Chloroacetamide (ClAA) addition, respectively. LC-MS analysis of the resulting peptide mixture was conducted on an Orbitrap Exploris 480 Mass Spectrometer (Thermo Fisher Scientific) directly coupled by ESI to an ACQUITY UPLC M-Class System (Waters) configured for 75-micron scale single-pump trapping. Peptides were separated on a nanoEase HSS C18 T3, 100A, 75 um x 250 mm analytical column (Waters, PN: 186008818) at a constant flow rate of 300 nl/min applying a piece-wise linear gradient from 5 to 32% solvent B in 45 min (solvent A: water incl. 0.1% formic acid, solvent B: acetonitrile incl. 0.1% formic acid). MS data acquisition was conducted in data-dependent mode. Data-dependent scans (MS2) were acquired in the Orbitrap mass analyzer at 30’000 resolution (norm. AGC target: 100%, maxIT: 119 ms) covering precursor ions in the scan range 350-1200 m/z and in charge states 2 to 7. Precursors were quadrupole isolated at 1.2 m/z and HCD fragmented at a NCE of 30. Precursor scans (MS1) were also conducted in the Orbitrap mass analyzer at 120’000 resolution (AGC target: 300%, maxIT: 50 ms) including lock mass correction. Resulting LC-MS data was processed using a scripted Philosopher workflow ^41^ followed by Label-Free Quantification using IonQuant ^42^. MS2 spectra were searched against the human reference proteome (UP000005640) by the MSFragger search engine 3.4, allowing for one missed tryptic cleavage and fixed carbamidomethylation of Cysteine, variable Methionine oxidation, and variable acetylation of the protein N-terminus after Methionine removal. Label-Free quantification was performed applying the FDR-controlled Match-Between-Runs algorithm as implemented by IonQuant 1.7.17. Differential abundance analysis was performed using proLFQua ^43^. Potential pray proteins were identified after robust scale (z-score) transformation, by filtered at a FDR cutoff of 5% and a fold-change of 2.

### Data Availability

Atomic coordinate files have been deposited in Protein Data Bank (PDB) under accession codes 9SPD (tRNA-LC) and 9SPE (tRNA-LC-PYROXD1 complex). Cryo-EM maps have been deposited in the Electron Microscopy Database (EMDB) under accession codes EMD-55078 (tRNA-LC-PYROXD1), EMD-55076

(refined map for tRNA-LC) and EMD-55075 (computationally enhanced map for tRNA-LC). Proteomics raw data and analysis scripts were submitted to PRIDE under project accession PXD068427.

## Supplementary Figures

**Figure S1.**
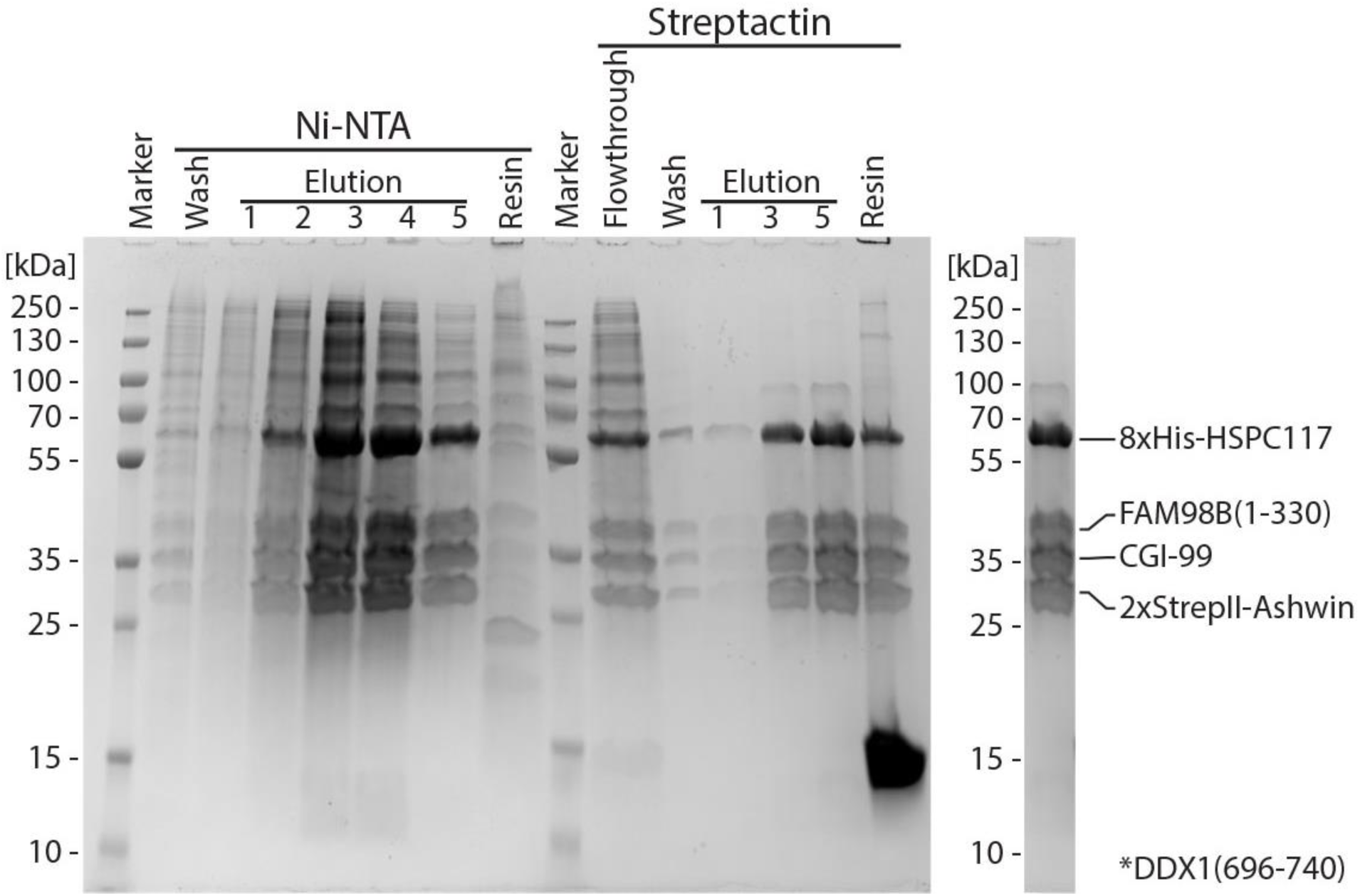
Purification of the tRNA-LC. Two-step purification of the tRNA-LC expressed in insect cells. In the first purification step recombinant tRNA-LC was purified by Ni-NTA affinity chromatography. The second purification step used Streptactin-affinity chromatography and desthiobiotin elution.

**Figure S2.**
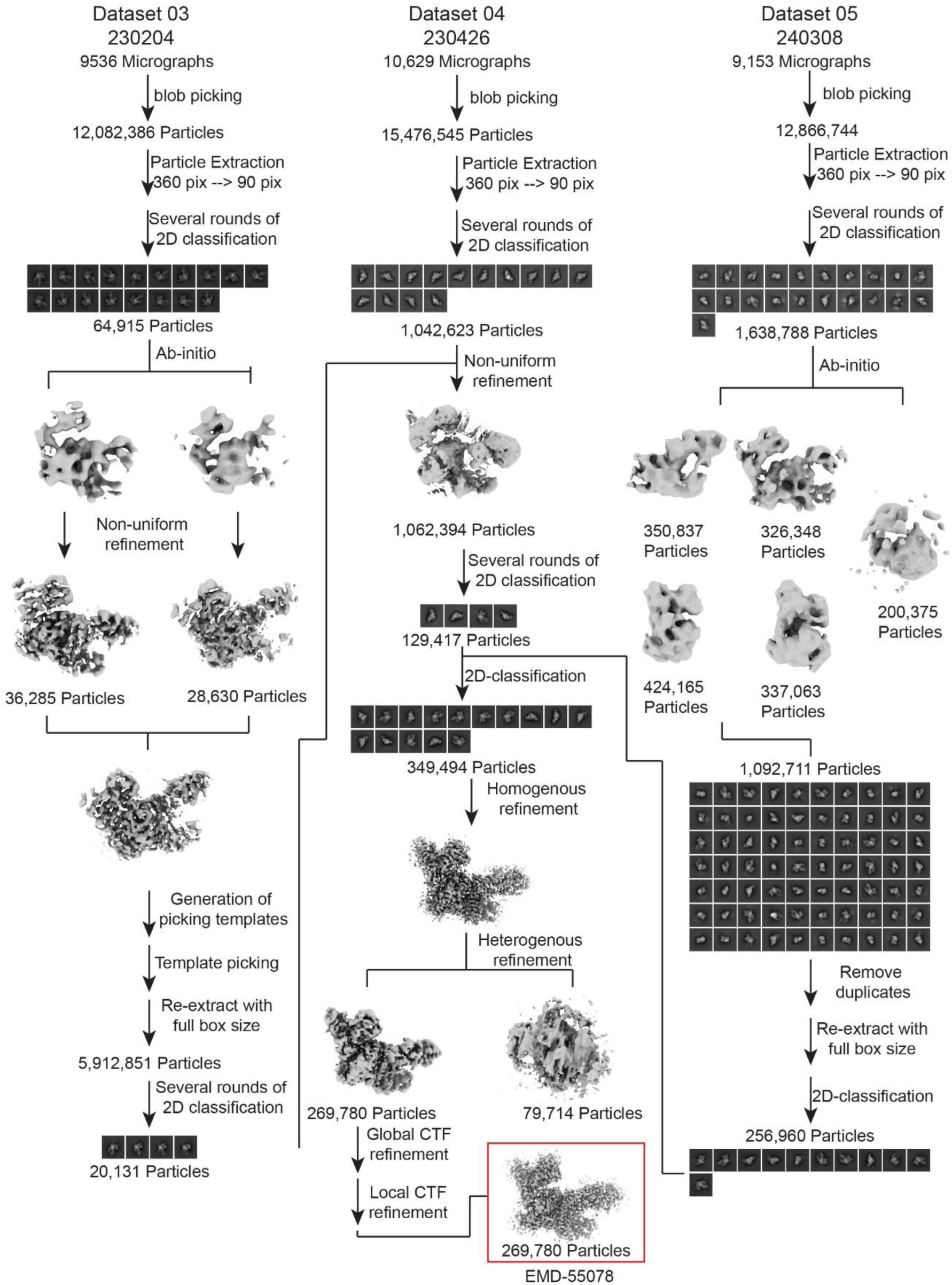
Cryo-EM data processing workflow for heteropentameric tRNA-LC. CryoSPARC processing tree for tRNA-LC.

**Figure S3.**
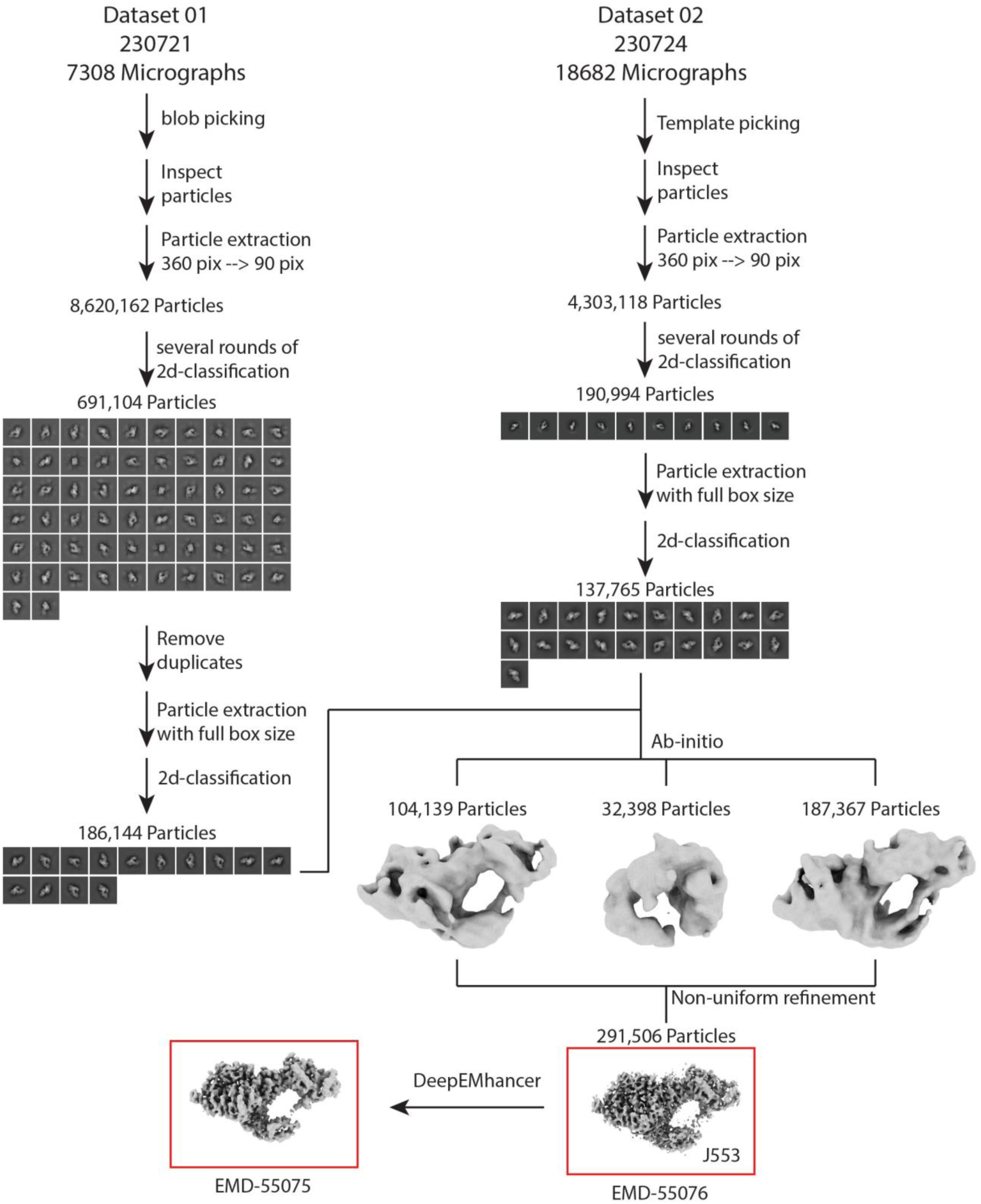
Cryo-EM data processing workflow for tRNA-LC complexed with PYROXD1. CryoSPARC processing tree for tRNA-LC complexed with PYROXD1.

**Figure S4.**
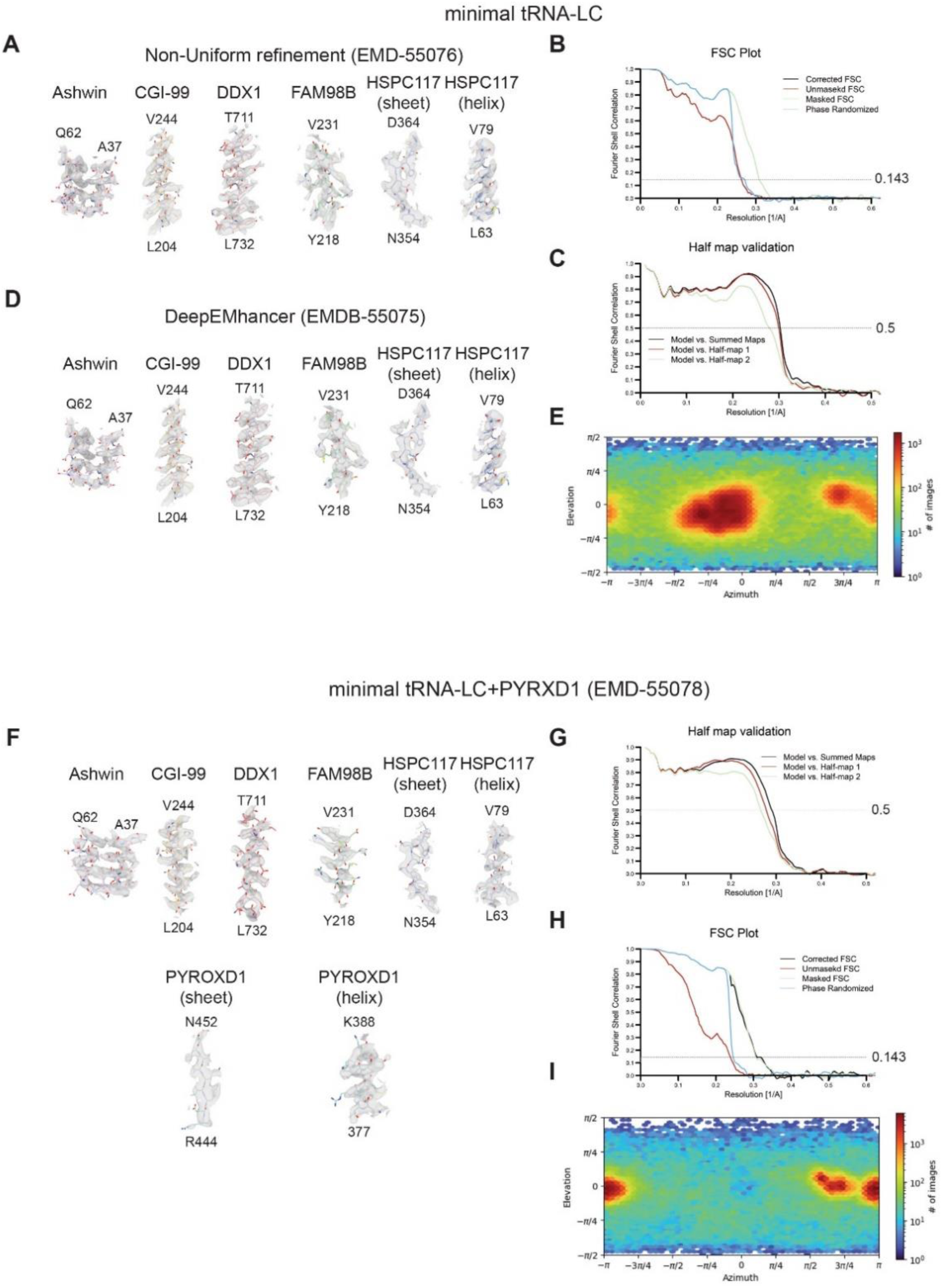
Cryo-EM model and map quality. (**A**) Representative helices and β-sheets modeled in corresponding cryo-EM densities for the pentameric tRNA-LC. (**B**) FSC-plot plot for tRNA-LC. (**C**) Model validation plots for the tRNA-LC. (**D**) Representative helices and β-sheets modelled in corresponding cryo-EM densities for the apo-tRNA after processing with DeepEMhancer. (**E**) Orientation distribution plot for the tRNA-LC. (**F**) Representative helices and β-sheets modelled in corresponding cryo-EM densities for tRNA-LC in complex with PYROXD1. (**G**) FSC-plot plot for tRNA-LC in complex with PYROXD1. (**H**) Model validation plots for tRNA-LC in complex with PYROXD1. (**I**) Orientation distribution plot for tRNA-LC in complex with PYROXD1.

**Figure S5.**
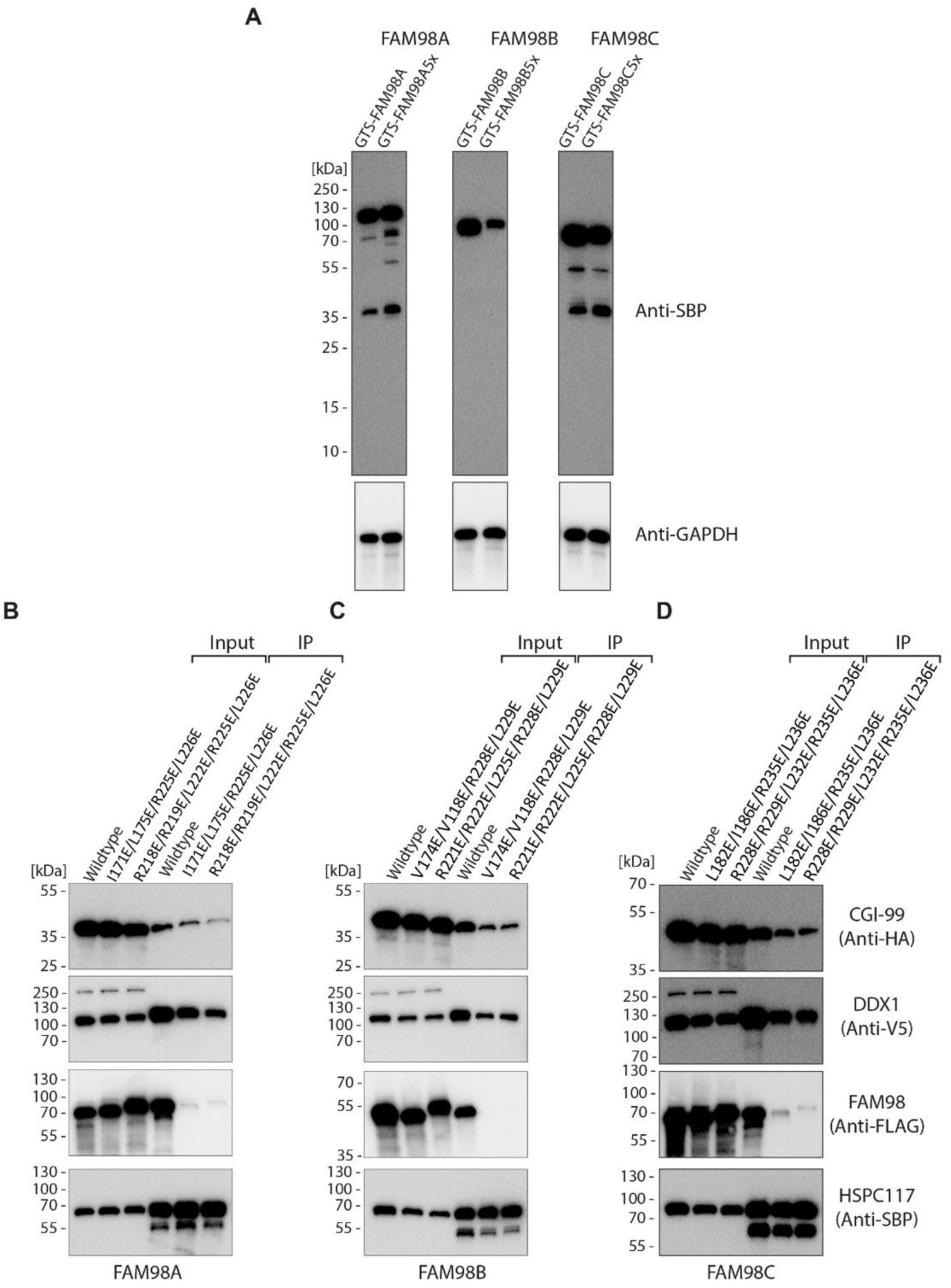
Generation and validation of FAM98 mutant cell lines. (**A**) Expression of mEGFP-TEV-SBP- (GTS-) tagged FAM98A-C paralogs in stably transfected *FAM98A-C* gene knock-out HEK293T cell lines, monitored by Western blot analysis using an anti-SBP antibody. 5x refers to mutant proteins containing five points mutations required to impair recruitment to the tRNA-LC: R218E/R219E/L222E/R225E/L226E^FAM98A^, R221E/R222E/L225E/R228E/L229E^FAM98B^ and R228E/R229E/L232E/R235E/L236E^FAM98C^. (**B,C,D**) Streptavidin affinity co-precipitation experiments to test structure-based mutations of FAM98A/B/C paralogs disrupting recruitment to the tRNA-LC.

**Supplementary Table S1.** Cryo-EM data collection, refinement, and validation statistics.

|  | tRNA-LC + PYROXD1 | tRNA-LC | tRNA-LC (deepEM) |
| --- | --- | --- | --- |
| **Data collection and processing** |  |  |  |
| Magnification | 130,000 | 130,000 | 130,000 |
| Voltage (kV) | 300 | 300 | 300 |
| Electron exposure (e<sup>-</sup>/Å<sup>2</sup>) | 50-76 | 60 | 60 |
| Defocus range (μm) | - 1.0 to -2.4 (-0.2 steps) | - 1.0 to -2.4 (-0.2 steps) | - 1.0 to -2.4 (-0.2 steps) |
| Pixel size (Å) | 0.65 | 0.65 | 0.65 |
| Symmetry imposed | C1 | C1 | C1 |
| Initial particle images (no.) | 29,197,719 | 12,923,250 | 12,923,250 |
| Final particle images (no.) | 269,780 | 291,506 | 291,506 |
| Map resolution (Å) | 3.23 | 3.25 | 3.25 |
| FSC threshold | 0.143 | 0.143 | 0.143 |
| Map resolution range (Å) | 3-5 | 3-5 | 3-5 |
| **Refinement** |  |  |  |
| Model resolution (Å) | 3.5 | 3.3 |  |
| FSC threshold | 0.5 | 0.5 |  |
| Map sharpening *B* factor (Å<sup>2</sup>) | 98.7 | 124.6 | 124.6 |
| Model composition |  |  |  |
| Non-hydrogen atoms | 11417 | 8130 |  |
| Protein residues | 1440 | 1039 |  |
| Nucleotides | 0 | 0 |  |
| Ligands | 6 | 0 |  |
| *B* factors (Å<sup>2</sup>) | min/max/mean | min/max/mean |  |
| Protein | 19.43/184.70/95.59 | 17.32/178.49/87.28 |  |
| Nucleotides | - | - |  |
| Ligands | 15.53/117.58/95.71 | - |  |
| R.m.s. deviations |  |  |  |
| Bond lengths (Å) | 0.003 | 0.002 |  |
| Bond angles (°) | 0.620 | 0.560 |  |
| Validation |  |  |  |
| MolProbity score | 1.86 | 1.38 |  |
| Clashscore | 8.69 | 5.11 |  |
| Poor rotamers (%) | 1.88 | 0.8 |  |
| Ramachandran plot |  |  |  |
| Favored (%) | 96.89 | 97.45 |  |
| Allowed (%) | 3.11 | 2.55 |  |
| Disallowed (%) | 0 | 0 |  |

## References

1. Goodman, H. M., Olson, M. V. & Hall, B. D. Nucleotide sequence of a mutant eukaryotic gene: the yeast tyrosine-inserting ochre suppressor SUP4-o. Proc. Natl. Acad. Sci. 74, 5453–5457 (1977).

2. Valenzuela, P., Venegas, A., Weinberg, F., Bishop, R. & Rutter, W. J. Structure of yeast phenylalanine-tRNA genes: an intervening DNA segment within the region coding for the tRNA. Proc. Natl. Acad. Sci. 75, 190–194 (1978).

3. Chan, P. P. & Lowe, T. M. GtRNAdb 2.0: an expanded database of transfer RNA genes identified in complete and draft genomes. Nucleic Acids Res. 44, D184–D189 (2016).

4. Gogakos, T. et al. Characterizing Expression and Processing of Precursor and Mature Human tRNAs by Hydro-tRNAseq and PAR-CLIP. Cell Rep. 20, 1463–1475 (2017).

5. Dantuluri, S., Schwer, B., Abdullahu, L., Damha, M. J. & Shuman, S. Activity and substrate specificity of *Candida*, *Aspergillus*, and *Coccidioides* Tpt1: essential tRNA splicing enzymes and potential antifungal targets. RNA 27, 616–627 (2021).

6. Popow, J., Schleiffer, A. & Martinez, J. Diversity and roles of (t)RNA ligases. Cell. Mol. Life Sci. 69, 2657–2670 (2012).

7. Fujishima, K. & Kanai, A. tRNA gene diversity in the three domains of life. Front. Genet. 5, (2014).

8. Popow, J. et al. HSPC117 Is the Essential Subunit of a Human tRNA Splicing Ligase Complex. Science 331, 760–764 (2011).

9. Popow, J., Jurkin, J., Schleiffer, A. & Martinez, J. Analysis of orthologous groups reveals archease and DDX1 as tRNA splicing factors. Nature 511, 104–107 (2014).

10. Banerjee, A., Ghosh, S., Goldgur, Y. & Shuman, S. Structure and two-metal mechanism of fungal tRNA ligase. Nucleic Acids Res. 47, 1428–1439 (2019).

11. Chakravarty, A. K., Subbotin, R., Chait, B. T. & Shuman, S. RNA ligase RtcB splices 3′-phosphate and 5′-OH ends via covalent RtcB-(histidinyl)-GMP and polynucleotide-(3′)pp(5′)G intermediates. Proc. Natl. Acad. Sci. 109, 6072–6077 (2012).

12. Greer, C. L., Peebles, C. L., Gegenheimer, P. & Abelson, J. Mechanism of action of a yeast RNA ligase in tRNA splicing. Cell 32, 537–546 (1983).

13. Kroupova, A. et al. Molecular architecture of the human tRNA ligase complex. eLife 10, e71656 (2021).

14. Akter, K. A., Mansour, M. A., Hyodo, T. & Senga, T. FAM98A associates with DDX1-C14orf166-FAM98B in a novel complex involved in colorectal cancer progression. Int. J. Biochem. Cell Biol. 84, 1–13 (2017).

15. Pazo, A. et al. hCLE/RTRAF-HSPC117-DDX1-FAM98B: A New Cap-Binding Complex That Activates mRNA Translation. Front. Physiol. 10, 92 (2019).

16. Pérez-González, A. et al. hCLE/C14orf166 Associates with DDX1-HSPC117-FAM98B in a Novel Transcription-Dependent Shuttling RNA-Transporting Complex. PLoS ONE 9, e90957 (2014).

17. Asanović, I. et al. The oxidoreductase PYROXD1 uses NAD(P)+ as an antioxidant to sustain tRNA ligase activity in pre-tRNA splicing and unfolded protein response. Mol. Cell 81, 2520–2532.e16 (2021).

18. Desai, K. K., Cheng, C. L., Bingman, C. A., Phillips, G. N. & Raines, R. T. A tRNA splicing operon: Archease endows RtcB with dual GTP/ATP cofactor specificity and accelerates RNA ligation. Nucleic Acids Res. 42, 3931–3942 (2014).

19. Gerber, J. L. et al. Structural and mechanistic insights into activation of the human RNA ligase RTCB by Archease. Nat. Commun. 15, 2378 (2024).

20. Loeff, L. et al. ar. Preprint at 10.1101/2023.04.06.535761 (2023).

21. O’Grady, G. L. et al. Variants in the Oxidoreductase PYROXD1 Cause Early-Onset Myopathy with Internalized Nuclei and Myofibrillar Disorganization. Am. J. Hum. Genet. 99, 1086–1105 (2016).

22. Schaffer, A. E. et al. CLP1 Founder Mutation Links tRNA Splicing and Maturation to Cerebellar Development and Neurodegeneration. Cell 157, 651–663 (2014).

23. Jurkin, J. et al. The mammalian tRNA ligase complex mediates splicing of *XBP1* mRNA and controls antibody secretion in plasma cells. EMBO J. 33, 2922–2936 (2014).

24. Kosmaczewski, S. G. et al. The R tc B RNA ligase is an essential component of the metazoan unfolded protein response. EMBO Rep. 15, 1278–1285 (2014).

25. Lu, Y., Liang, F.-X. & Wang, X. A Synthetic Biology Approach Identifies the Mammalian UPR RNA Ligase RtcB. Mol. Cell 55, 758–770 (2014).

26. Yanagitani, K., Kimata, Y., Kadokura, H. & Kohno, K. Translational Pausing Ensures Membrane Targeting and Cytoplasmic Splicing of *XBP1u* mRNA. Science 331, 586–589 (2011).

27. Edgcomb, S. P. et al. DDX1 Is an RNA-Dependent ATPase Involved in HIV-1 Rev Function and Virus Replication. J. Mol. Biol. 415, 61–74 (2012).

28. Zhang, Z. et al. DDX1, DDX21, and DHX36 Helicases Form a Complex with the Adaptor Molecule TRIF to Sense dsRNA in Dendritic Cells. Immunity 34, 866–878 (2011).

29. Jumper, J. et al. Highly accurate protein structure prediction with AlphaFold. Nature 596, 583–589 (2021).

30. Suzuki, T. et al. DDX1 is required for non-spliceosomal splicing of tRNAs but not of XBP1 mRNA. *Commun*. Biol. 8, 92 (2025).

31. Nemudraia, A., Nemudryi, A. & Wiedenheft, B. Repair of CRISPR-guided RNA breaks enables site-specific RNA excision in human cells. Science 384, 808–814 (2024).

32. Loeff, L. et al. Mechanistic basis for PYROXD1-mediated protection of the human tRNA ligase complex against oxidative inactivation. Nat. Struct. Mol. Biol. 32, 1205–1212 (2025).

33. Chamera, S. et al. Structural and biochemical characterization of the 3′-5′ tRNA splicing ligases. J. Biol. Chem. 301, 108506 (2025).

34. Yang, J. et al. Polyglycine-mediated aggregation of FAM98B disrupts tRNA processing in GGC repeat disorders. Science 389, eado2403 (2025).

35. Patil, S. S. et al. Novel gene *ashwin* functions in *Xenopus* cell survival and anteroposterior patterning. Dev. Dyn. 235, 1895–1907 (2006).

36. Leitner, M. et al. Ashwin and FAM98 paralogs define nuclear and cytoplasmic RNA ligase complexes for tRNA biogenesis and the unfolded protein response. Preprint at 10.1101/2025.08.01.668163 (2025).

37. Weissmann, F. et al. biGBac enables rapid gene assembly for the expression of large multisubunit protein complexes. Proc. Natl. Acad. Sci. 113, E2564–E2569 (2016).

38. Berger, I., Fitzgerald, D. J. & Richmond, T. J. Baculovirus expression system for heterologous multiprotein complexes. Nat. Biotechnol. 22, 1583–1587 (2004).

39. Zhang, K. Gctf: Real-time CTF determination and correction. J. Struct. Biol. 193, 1–12 (2016).

40. Sanchez-Garcia, R. et al. DeepEMhancer: a deep learning solution for cryo-EM volume post-processing. *Commun*. Biol. 4, 874 (2021).

41. Da Veiga Leprevost, F. et al. Philosopher: a versatile toolkit for shotgun proteomics data analysis. Nat. Methods 17, 869–870 (2020).

42. Yu, F., Haynes, S. E. & Nesvizhskii, A. I. IonQuant Enables Accurate and Sensitive Label-Free Quantification With FDR-Controlled Match-Between-Runs. Mol. Cell. Proteomics MCP 20, 100077 (2021).

43. Wolski, W. E. et al. *prolfqua* : A Comprehensive *R* -Package for Proteomics Differential Expression Analysis. J. Proteome Res. 22, 1092–1104 (2023).

